# Conditioning dentinal collagen by glycation improves long-term bond strength

**DOI:** 10.64898/2026.01.23.701305

**Authors:** Wafa Alzubaidi, Bowen Wang, Mina Vaez, Mehrnoosh Neshatian, Sebastian Aguayo, Steven Thorpe, Eszter Somogyi-Ganss, Laurent Bozec

**Affiliations:** Faculty of Dentistry, University of Toronto, Toronto, ON, Canada; Department of Material Science and Engineering, University of Toronto. Toronto, ON; Dentistry School, Faculty of Medicine, Pontificia Universidad Católica de Chile, Santiago, Chile; Institute for Biological and Medical Engineering, Schools of Engineering, Medicine and Biological Sciences, Pontificia Universidad Católica de Chile, Santiago, Chile

**Keywords:** dentinal collagen, collagen crosslinking, methylglyoxal, ATR-FTIR, enzymatic degradation, modulus of elasticity

## Abstract

**Objectives:** This study examined methylglyoxal (MGO) as a collagen crosslinker to reinforce demineralized dentin, enhance its enzymatic resistance, and improve long-term durability of the resin–dentin bond.

**Methods:** Demineralized dentinal collagen films were treated with 0.5, 1, or 3 M MGO and subsequently exposed to collagenase to assess their resistance to enzymatic degradation. MGO-induced crosslink formation was monitored using Attenuated Total Reflectance–Fourier Transform Infrared spectroscopy (ATR-FTIR) spectroscopy by tracking the carbohydrate-associated band at 1180 cm⁻¹. The apparent elastic modulus of the treated specimens was measured using a three-point bending test. For bond strength evaluation, resin–dentin beams were prepared and tested using microtensile bond strength (μTBS) to assess the influence of MGO pretreatment on interfacial adhesion.

**Results:** The ATR-FTIR spectra demonstrated increased intensity in the carbohydrate double bands (1000–1180 cm⁻¹) in glycated samples compared to the control. Glycation with 3 M MGO exhibited the highest resistance to enzymatic degradation, persisting for up to 60 hours with a 78-fold increase in resistance factor compared to the control group (p < 0.05). Furthermore, glycation with 3 M MGO resulted in a 4-fold increase in elastic modulus compared with the control group. Notably, the functionalized dentin retained its improved mechanical properties even after collagenase exposure, whereas the control group experienced a significant 68.2% reduction in elastic modulus (p = 0.002). While MGO pretreatment did not influence resin infiltration or initial μTBS (≈30–35 MPa), it maintained its original bond strength after one month of collagenase challenge. In contrast, the control group exhibited a significant reduction, decreasing to 17 ± 5.5 MPa compared with its initial value (p < 0.01).

**Conclusion:** MGO demonstrated efficacy in enhancing the mechanical properties and enzymatic stability of collagen as well as improving the resistance of the bonded interface to enzymatic degradation.

**Significance:** MGO pretreatment maintains the long-term stability of the resin–dentin interface by protecting dentinal collagen from enzymatic degradation, without compromising initial bonding performance.

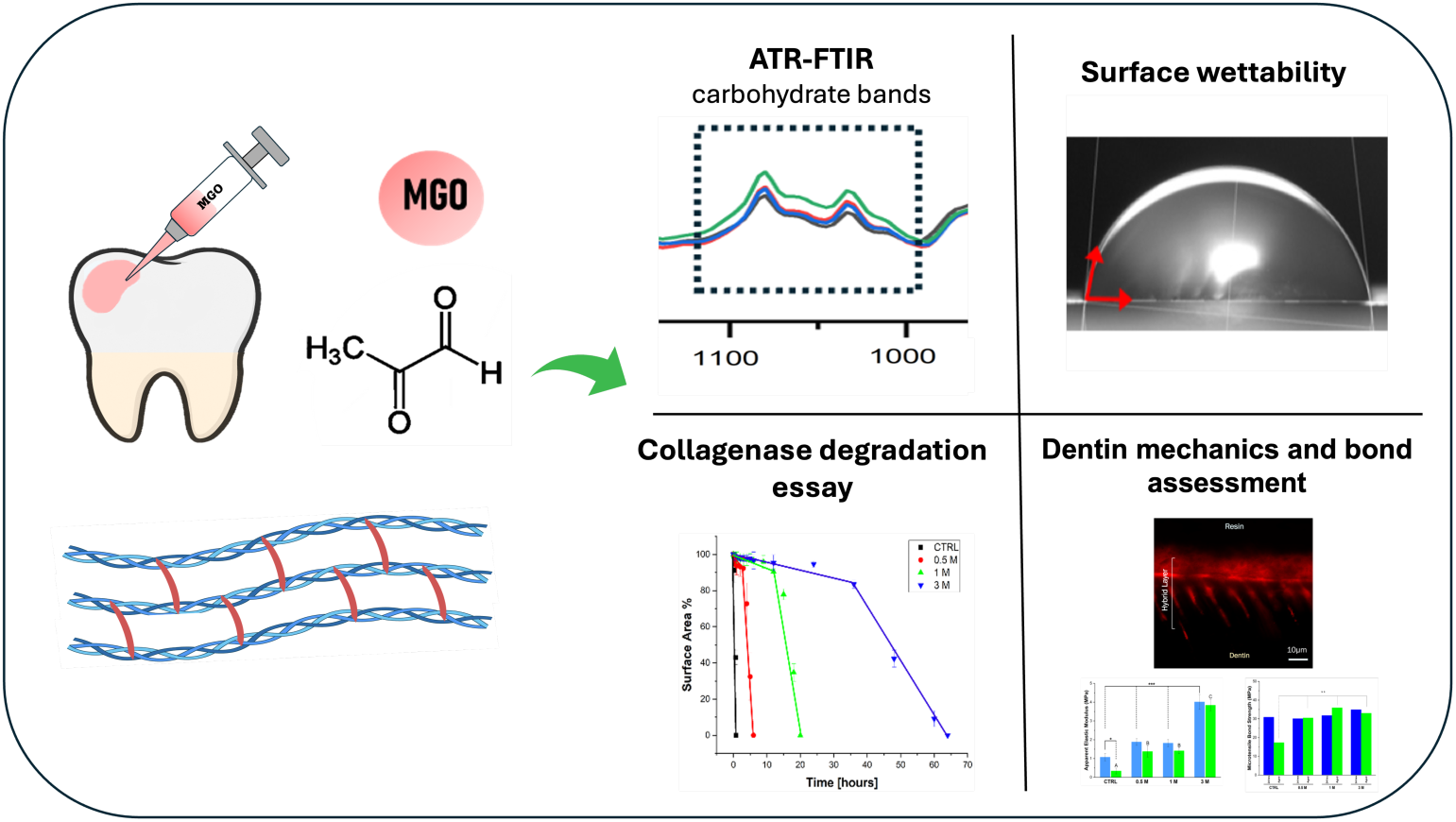

## 1 Introduction

Composite resin restorations have evolved the field of restorative dentistry by combining aesthetic appeal with satisfactory mechanical performance[1, 2]. However, ensuring a long-lasting bond between resin restorations and dentin remains a challenge, impacting clinical success and increasing the economic burden[3, 4]. Resin restorations, when bonded to dentin, encounter various challenges, including mechanical stress, chemical degradation, and biodegradation from bacteria or host-derived enzymes, all of which undermine the longevity of the adhesive interface[5]. Bonding to dentin is intrinsically more difficult than bonding to enamel due to its higher water and organic composition, featuring an intricate network of tubules within a collagen-rich matrix that complicates adhesive stability[6]. This bond is established through chemical interactions[7] and micromechanical interlocking between the resin material and the demineralized surface[8]. The clinical protocol typically begins with acid etching, which removes the smear layer, exposes collagen fibrils, and prepares the substrate for resin monomer infiltration, thereby forming the hybrid layer[8]. However, excessive demineralization can compromise collagen’s structural integrity, rendering it more susceptible to hydrolysis and enzymatic degradation. Phosphoric acid breaks down intermolecular crosslinks, causing the unsupported dentinal collagen network to collapse[9]. This structural collapse diminishes resin infiltration efficiency [10] and heightens the hybrid layer’s susceptibility to degradation by endogenous matrix metalloproteinases (MMPs) [11]. MMP-8 targets type I collagen, and gelatinase (MMP-2 and MMP-9) are also responsible for organic hydrolysis in demineralized dentin, altering the surface morphology[5]. Furthermore, leached bisGMA, TEGDMA, and their by-products enhance MMP-8 and -9 activity[12]. Finally, dentin etching creates an acidic environment that activates MMPs, degrading collagen molecules and potentially weakening bond integrity between resin and dentin[13]. The successful bonding of resin restorations to dentin relies on establishing robust adhesion with the collagen fibrils embedded within the dentin matrix[14]. Therefore, collagen crosslinking has been explored to improve the matrix’s mechanical properties while enhancing its resistance to enzymatic degradation. Exogenous crosslinking agents, such as proanthocyanidins[15, 16], glutaraldehyde[17], lignin[18], and riboflavin[19], have shown promise in improving resin-dentin bond strength. Recently, methylglyoxal (MGO), a naturally occurring sugar-derived crosslinker, has gained attention for enhancing collagen-based tissues through non-enzymatic crosslinking, forming advanced glycation end-products (AGEs)[20, 21]. A study by Vaez et al. in 2023 demonstrated improved collagen fibril formation, increased scaffold stiffness, and enhanced resistance to proteolytic degradation in glycated type-I collagen scaffolds compared to the control [22].

Enhancing the structural integrity of exposed collagen is increasingly recognized as a promising strategy to promote durable adhesion at the biomaterial–dentin interface. This study aims to evaluate the effects of controlled glycation using methylglyoxal (MGO) on the structural and mechanical properties of demineralized dentinal collagen, thereby reinforcing the collagen matrix against enzymatic degradation and enhancing its potential to support long-term resin adhesive performance.

## 2 Materials and Methods

The University of Toronto Research Ethics Board (REB 44300) approved the collection of 39 extracted human permanent third molars. The teeth were scaled to remove soft tissue, stored in a 0.1% sodium azide solution, and refrigerated at 4°C until use. All mineralized dentin sections were prepared using a water-cooled low-speed diamond blade (Buehler Ltd., Lake Bluff, IL, USA) mounted on an Isomet low-speed saw (Buehler, Lake Bluff, IL, USA). Figure 1 provides an overview of the experimental workflow.

**Figure 1.**
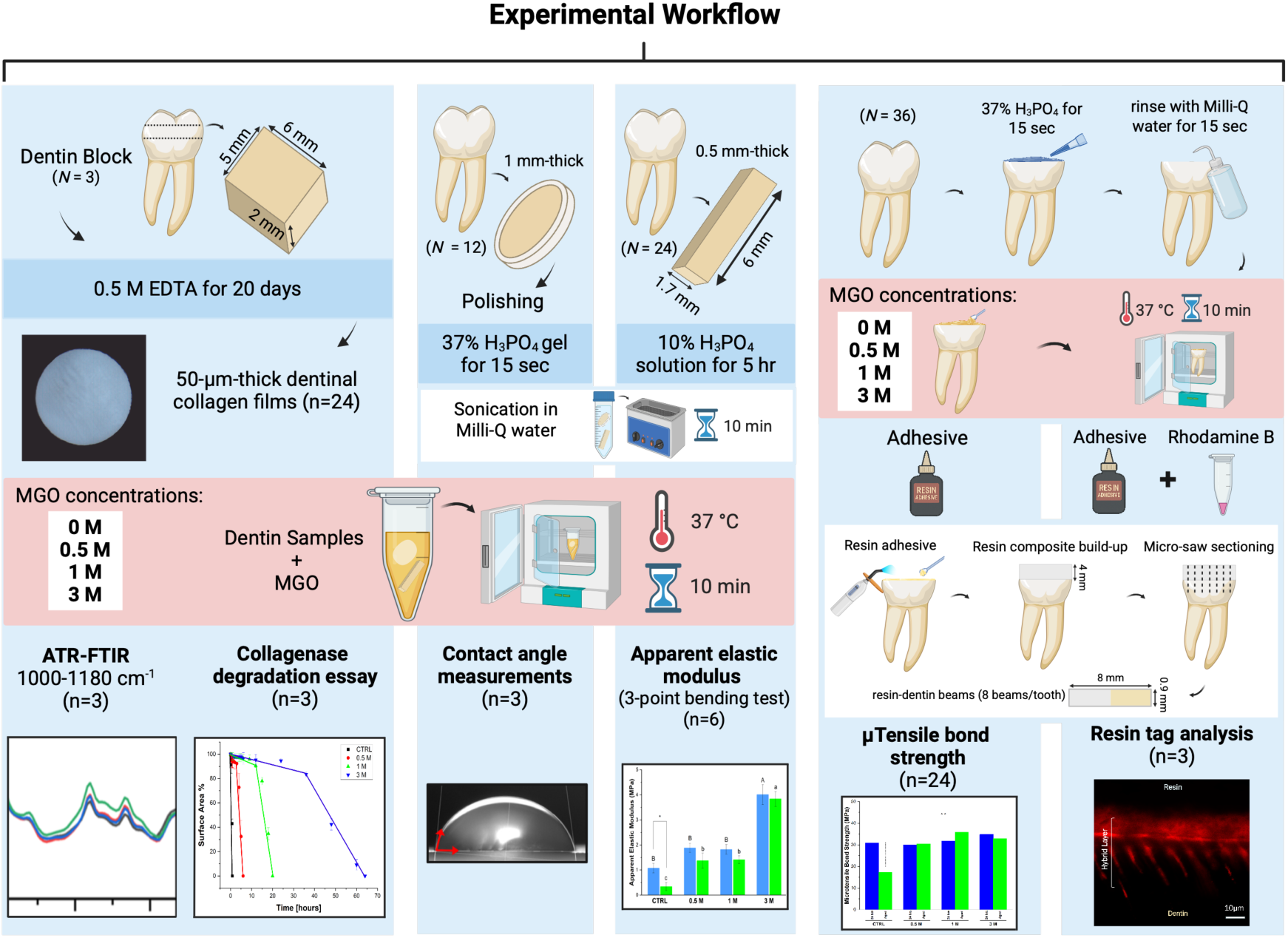
Workflow Chart for the Study: The capital letter N refers to the total number of teeth. The small letter n refers to the number of samples per group randomly assigned to each experiment. In the functionalization protocol, each experiment included 4 groups (control (0 M MGO) and 3 concentrations of MGO (0.5 M, 1 M, and 3 M). EDTA (ethylenediaminetetraacetic acid), H₃PO₄ (phosphoric acid). *C.H.C; Clostridium Histolyticum* Collagenase.

For Attenuated Total Reflectance - Fourier Transform Infrared Spectroscopy (ATR-FTIR) and collagenase degradation assessment, three coronal dentin blocks (6 x 5 x 2 mm) were sectioned and demineralized for 20 days in 0.5 M EDTA. After complete demineralization, 24 (50 µm thick) sections were prepared using a microtome (Leica CM1850, Deerfield, IL, USA). A 4-mm hole punch was performed on each section to standardize the surface area. Samples were kept in PBS at 4°C before being randomly distributed for the subsequent testing procedure. For collagen glycation, samples were immersed in MGO (0.5 M, 1 M, and 3 M) for 10 minutes at 37°C. Afterwards, they were removed and washed extensively with ultrapure water for 10 seconds. The samples were immediately used for subsequent testing.

### 2.1 Confirming the formation of MGO-derived crosslinks

ATR-FTIR monitored the intensity of the carbohydrate double band between 1000 and 1180 cm^-1^. Spectra were baseline-corrected and normalized before plotting in OriginLab software (OriginLab Corp., Northampton, MA, USA).

### 2.2 Assessing the resistance to complete degradation

Collagenase *Clostridium histolyticum* (CCH) (Sigma-Aldrich, MA, United States) was prepared by diluting the powder in 50 mM Tris-HCl buffer (dissolved in ultrapure water and adjusted to pH 7.4 with HCl). Films were incubated in a 1 ml solution containing 5 mg/ml of collagenase at 37°C until complete degradation occurred (n = 3). Samples were digitally imaged at set time points using a stereomicroscope (Nikon, SMZ800, Canada). The percentage of surface area during degradation was calculated using Image J (version 1.54) with a contour analysis routine and plotted as a function of time.

### 2.3 Surface wettability assessment

Twelve teeth were sectioned into 12 coronal dentin discs (1 ± 0.2 mm thick). Disks were polished with silicone-carbide grit papers (grades 500, 1200, 2400). Samples were sonicated in DI water for 10 minutes, then demineralized in 37% phosphoric acid gel for 15 seconds and rinsed thoroughly with UP water for 10 seconds. Finally, samples were treated with a micro-brush with MGO: 0.5 M, 1 M, and 3 M for 10 minutes at 37°C; samples without treatment were used as control. The samples were mounted on an XYZ-tilting stage, and the contact angles were measured using a 5 µL DI water drop. The water drop was captured at 15 seconds using a FLIR Blackfly Monochrome BFS-U3 high-speed camera. The right and left contact angles were calculated using ImageJ to obtain a mean θ value.

### 2.4 Testing the mechanical performance of functionalized dentin

Twenty-four dentin beams (0.5 × 1.7 × 6 mm) were sectioned, demineralized in 10 % phosphoric acid for 5 hours based on a previous protocol[23], sonicated in ultrapure water for 10 minutes, and stored in PBS. Samples were assigned to control or 3 MGO-treated groups (n = 6). Mechanical testing was done using a three-point bending setup on the ElectroPuls® E8501 (Instron Inc., Canton, USA). The apparent modulus (AEM) was measured 24 hours post-glycation by loading samples to maximum bending at 1 mm/min crosshead speed. After a 6-hour challenge in 0.1 mg/ml collagenase solution, the modulus was remeasured under identical conditions. Linear regression determined the steepest slope of each force-displacement curve. The supported span length was 2 mm. The apparent elastic modulus was calculated using the formula:

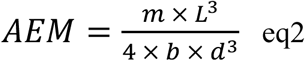

Where: **m** = slope (N/mm); **L** = support span (mm); **d** = thickness of beam (mm); **b** = width of beam (mm)

### 2.5 Testing the resin-dentin bond performance

Twenty-four teeth were randomly assigned to four experimental groups (n = 6 per group): (1) CTRL (no pretreatment); (2) 0.5 M MGO; (3) 1 M MGO; and (4) 3 M MGO.

#### Adhesive bonding protocol

All specimens were treated using a standardized etch-and-rinse approach. Dentin was etched with 37% phosphoric acid gel (Scotchbond™ Etchant, 3M) for 15 s, rinsed for 15 s, and kept visibly moist without pooling. Following phosphoric acid etching, each solution was applied to the exposed dentin surface for 10 min at 37 °C, rinsed for 30 s with distilled water, and gently blot-dried to maintain a moist surface. Adper™ Single Bond 2 (3M ESPE, USA) was applied in two consecutive coats with active agitation for 15 s, air-thinned for 5 s, and light-cured for 20 s using an LED unit (550 mW/cm²). Composite build-up was performed using Filtek™ One Bulk Fill (3M) in increments to a height of ∼4 mm, with each increment light-cured for 40 s.

After 24 h of water storage at 37 °C, each tooth was sectioned longitudinally under water cooling using a low-speed diamond saw to obtain resin–dentin sticks (∼0.9 × 0.9 mm²). Eight sticks were prepared per tooth. Four sticks were allocated to the immediate group and stored in PBS at 37 °C for 24 h, whereas the remaining four sticks comprised the aged group and were stored in 0.1% collagenase (Type I, Sigma-Aldrich, USA) at 37 °C for 30 days. The collagenase solution was refreshed every three days to maintain enzymatic activity.

For microtensile bond strength (µTBS) testing, each stick was secured to a microtensile jig using cyanoacrylate and aligned along its longitudinal axis. Tensile testing was performed using an ElectroPuls® E1000 testing machine (Instron Inc., USA) at a crosshead speed of 0.5 mm/min until failure. The bonded cross-sectional area was measured after debonding using a calibrated digital micrometre, and µTBS values (MPa) were calculated by dividing the maximum load at failure (N) by the measured bonded area (mm²).

For resin tag evaluation, three teeth per group were bonded using an adhesive doped with 0.1 wt% Rhodamine B (Sigma-Aldrich, USA)[24]. Teeth were processed using the same bonding and sectioning protocol. The resin–dentin interface was visualized using a Zeiss Axio Observer 7 inverted confocal laser scanning microscope equipped with an LSM 800 scan head and ZEN software (Carl Zeiss, Germany), using a 63× oil-immersion objective (numerical aperture = 1.4). Fluorescence emission from Rhodamine B was recorded in the 570–590 nm range. Serial optical sections were acquired along a fixed axis to a depth of 20 µm, with a z-step size of 0.3 µm. For each specimen, ten interfacial images were randomly captured from a single slab[24]. Resin tag length was measured using ImageJ software.

### 2.5 Statistical analysis

Before statistical analysis, the assumptions of parametric testing were evaluated. Normality of data distribution was assessed using the Shapiro–Wilk test, and homogeneity of variances was verified using Levene’s test. Upon confirmation, one-way ANOVA followed by Tukey’s post hoc test was performed to determine statistical significance. All analyses were conducted using OriginLab software (OriginLab Corp., Northampton, MA, USA), with a significance level set at p < 0.05.

## 3 Results and Discussions

### 3.1 Confirming the formation of MGO-derived cross-links & surface wettability assessment

Having demonstrated how etching can significantly affect the properties of dentinal collagen, it is appropriate to explore avenues that may either restore the collagen’s internal crosslinks lost due to etching or supplement the native dentin with additional crosslinks before etching. To test this, we functionalized the samples with 0.5 M, 1 M, and 3 M MGO solutions (the defined concentrations were based on previous research [25]). The biomolecular fingerprint of MGO-derived crosslinks in ATR-FTIR consists of carbohydrate double bands in the region between 1000-1180 cm^-1^ (Figure 2 A). These bands arise from the vibrations of C-O and C-C sugar moieties in AGEs. The FTIR spectra of the 3 M glycated group exhibited the strongest intensity among the other groups, particularly when compared to the control. This increased intensity in the carbohydrate double band confirms the formation of AGEs within the collagen films. A similar observation was reported in a study by Tiedemann et al. 2022[25], where treating the root dentin of a human tooth with 3 M MGO significantly increased sugar band intensity compared to the control. The spectra exhibit a slight enhancement at concentrations of 0.5 M and 1 M, compared to the control group, with no substantial change in intensity observed between these concentrations.

**Figure 2.**
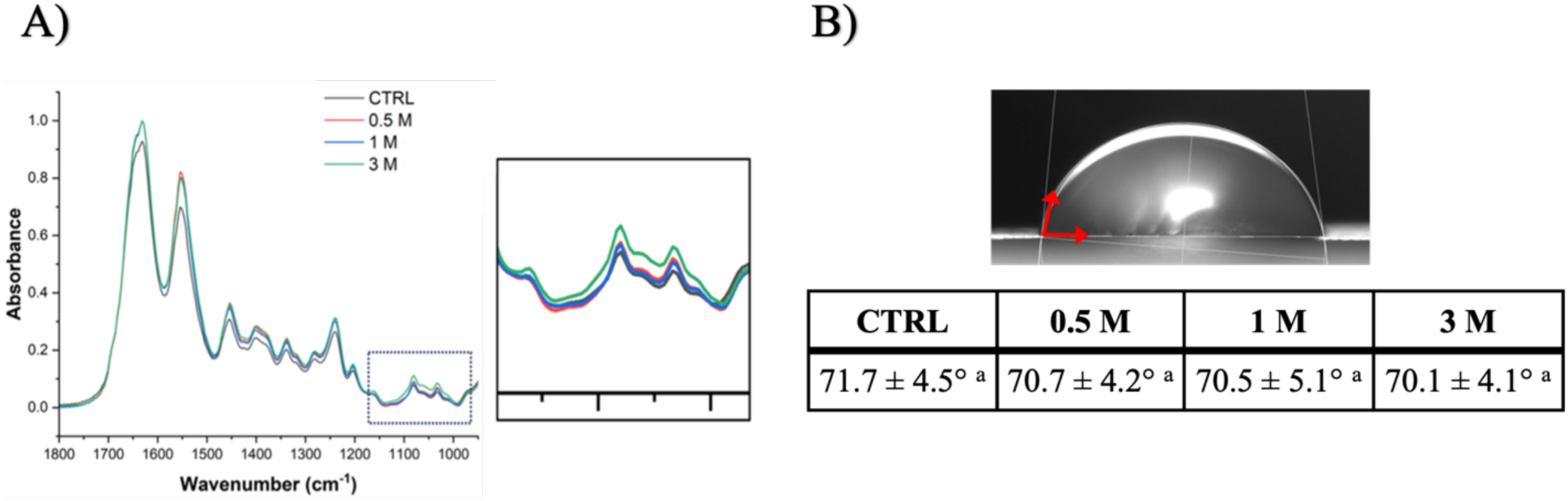
A) Normalized and baseline corrected ATR-FTIR spectra of control (black line), and glycated collagen incubated with 0.5 M, 1 M, and 3 M MGO (coloured lines). **B)** Mean and SD (n=3) of contact angle measurements for the control and 3 concentrations of MGO (p < 0.05).

Surface wettability refers to the extent to which a liquid spreads on a solid surface, determining whether the surface is hydrophilic (contact angle < 90°) or hydrophobic (contact angle > 90°)[26]. It is influenced by factors such as surface energy, roughness, and hydration status[26–28] and is typically assessed using the contact angle of a liquid droplet, often deionized (DI) water. Demineralized dentin is classified as hydrophilic, with contact angles of DI water drops ranging from 25° to 70°, depending on the time at which it was recorded[29–32]. The experiment (Figure 2 B) demonstrated that the mean contact angle for all groups ranged from 70.1° to 71.1°, with no statistically significant differences observed between the MGO-treated groups and the control (p > 0.05). This result suggests comparable surface wettability across all groups. While previous research has examined the effects of MGO on surface potential at the molecular level[33], studies explicitly focusing on its impact on surface wettability remain scarce. The similarity in contact angle measurements between the glycated and control samples implies that MGO treatment does not alter dentin surface wettability. This indicates that priming dentin with MGO before bonding is unlikely to interfere with the spreading behavior of adhesive monomers.

### 3.2 Enzymatic degradation susceptibility

Control and glycated collagen samples were exposed to a collagenase solution, and their resistance to complete biodegradation was monitored using time-lapse digital imaging. The control group exhibited the lowest resistance, with complete degradation occurring within 50 minutes (Figure 3 A). Increasing MGO concentrations significantly extended the time required for complete degradation. Collagen functionalized with 0.5 M and 1 M MGO degraded after approximately 6 and 20 hours, respectively, while the 3 M MGO glycation exhibited the highest resistance, lasting up to 60 hours. The percentage of the surface area is plotted as a function of time for all groups (Figure 3 B). The surface area plot shows that collagen films undergo a slow initial deterioration phase during enzymatic degradation, followed by rapid degradation (Figure 3 B). During the slow degradation phase, collagenase attacks non-specific sites in the triple helix structure of collagen molecules, disrupting these triple helices. As the degradation progresses, the breakdown of the triple helices starts to unwind and lose their stable conformation. This unwinding exposes more internal peptide bonds within the collagen molecules. The exposed peptide bonds within collagen are susceptible to more enzymatic cleavage or hydrolysis, leading to fragmentation of collagen molecules and significant loss of fibril integrity and functionality[34, 35].

**Figure 3.**
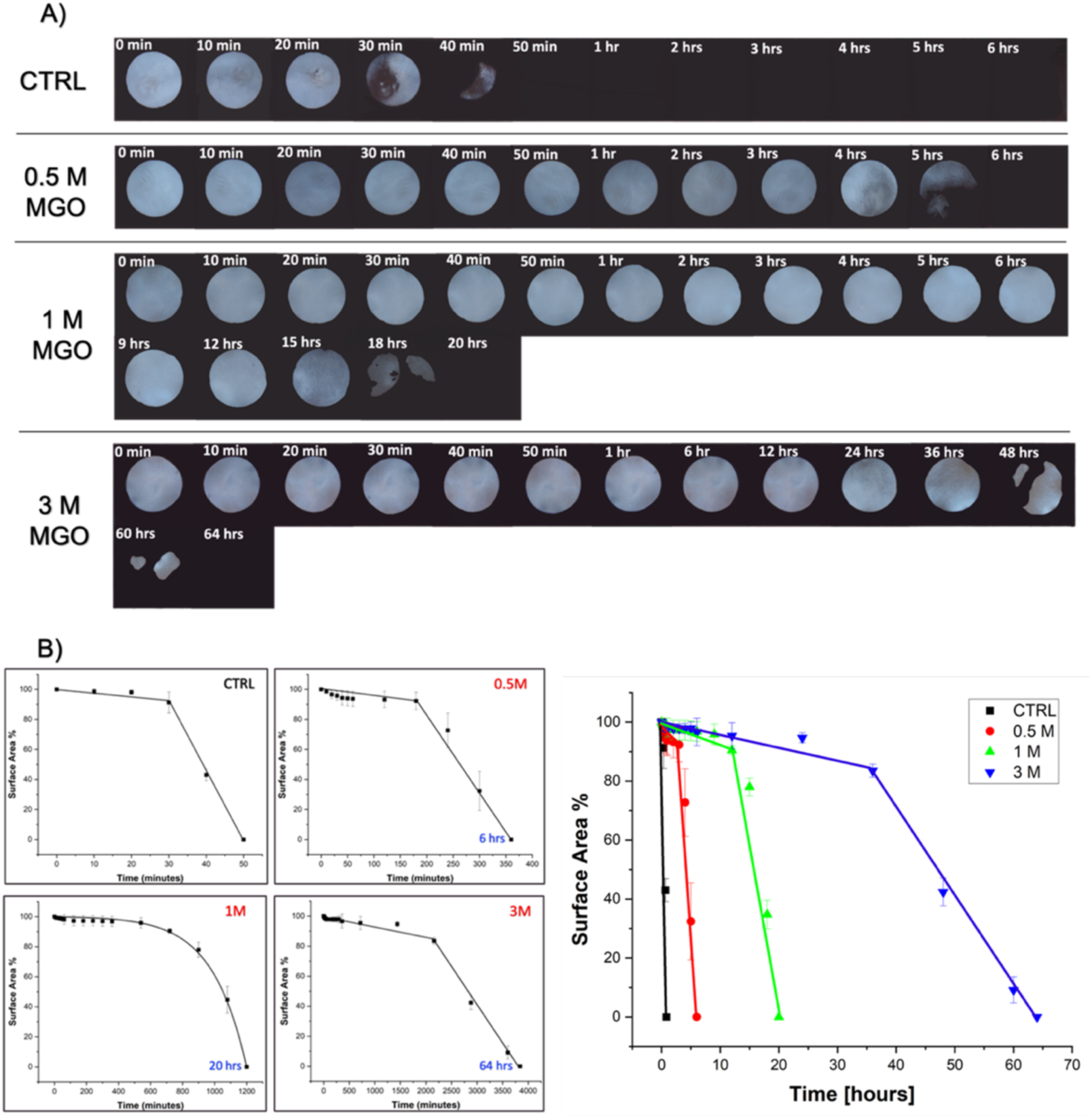
A) Time-lapse digital imaging of the control and functionalized demineralized dentinal films. Time measurements are expressed as “min” for minutes and “hrs” for hours. **B)** Representative plots of the progressive reduction in surface area caused by bacterial degradation as a function of time.

**Figure 4.**
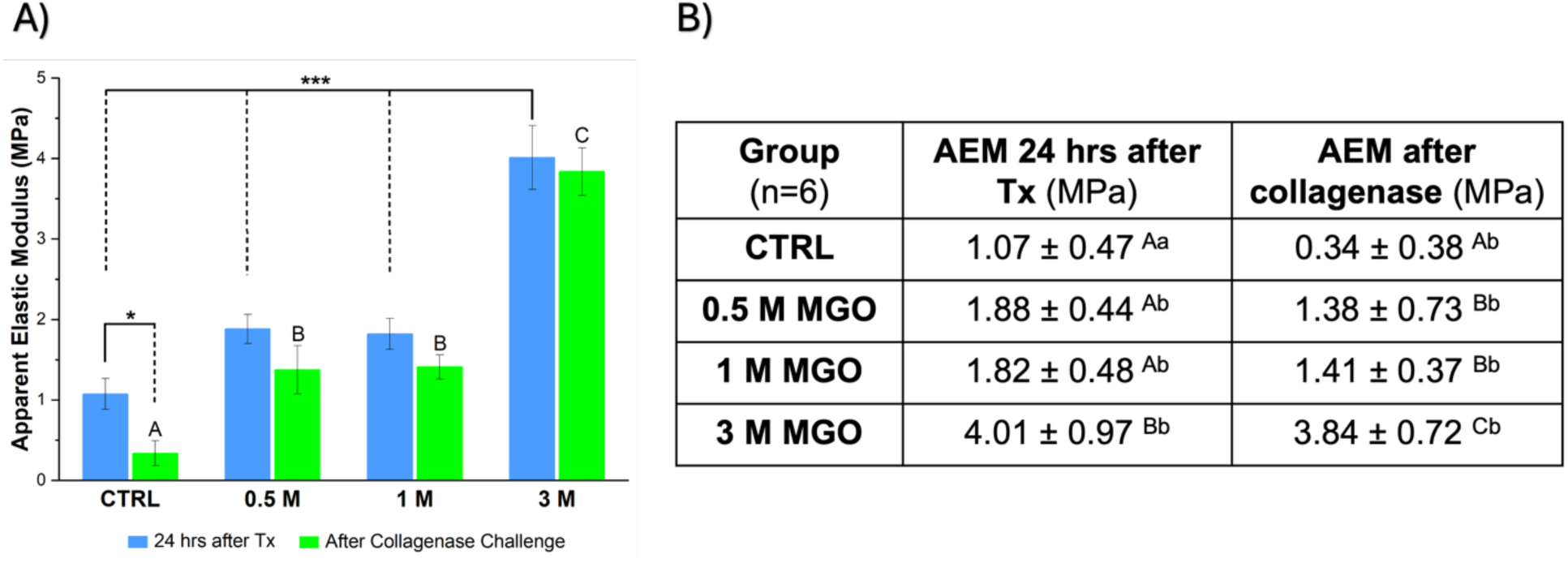
Apparent elastic modulus (AEM) of control and methylglyoxal (MGO)-treated samples before and after collagenase challenge. **(A)** The bar graph illustrates AEM values for untreated (CTRL) samples and samples treated with three different MGO concentrations (0.5 M, 1 M, and 3 M). Blue bars represent AEM values measured 24 hours post-treatment (Tx), while green bars represent AEM values after 6 hours of collagenase degradation. Statistical significance between groups post-treatment is indicated by asterisks (* p < 0.05, *** p < 0.001). Differences between groups after the collagenase challenge are denoted by uppercase letters (p < 0.05). **(B)** The table summarizes AEM values (Mean ± SD) for each group. Uppercase letters indicate differences across columns (between groups), while lowercase letters represent differences within rows (within groups), p < 0.05.

The onset of degradation was calculated for each group (Table 1). Control degraded at 28 ± 2 mins. With 0.5 M MGO, the onset of degradation was delayed to 3.4 ± 0.5 hours, demonstrating MGO’s stabilizing effect. At 1 M MGO, onset was 13.50 ± 1.5 hrs (p < 0.05), and at 3 M MGO, it was 35.9 ± 0.6 hrs, indicating stronger crosslinking. The resistance factor increased 7.2-fold and 24-fold for 0.5 M and 1 M MGO and 78-fold for 3 M MGO (p < 0.05). This concludes that higher MGO concentrations enhance collagen stability and its resistance to enzymatic degradation. A similar study found that lower MGO concentrations (up to 100 mM) slowed and delayed degradation in glycated collagen type I scaffolds[22].

**Table 1.** The onset of degradation (OD), time to complete degradation and resistance factor are presented as mean ± SD (n=3). Statistical significance is indicated by different letters in the resistance factor (p < 0.05).

| **Group**<br>(n=3) | **Onset of Degradation (OD)** | **Time to Complete Degradation** | **Resistance Factor**<br>($p < 0.05$) |
| --- | --- | --- | --- |
| **Control** | ($28 \pm 2$) mins | ($48 \pm 2$) mins | 1.0 <sup>a</sup> |
| **0.5 M MGO** | ($3.4 \pm 0.5$) hrs | ($5.8 \pm 0.3$) hrs | 7.2 <sup>b</sup> |
| **1 M MGO** | ($13.5 \pm 1.5$) hrs | ($20 \pm 2$) hrs | 24 <sup>c</sup> |
| **3 M MGO** | ($35.9 \pm 0.6$) hrs | ($63.3 \pm 4.5$) hrs | 78 <sup>d</sup> |

### 3.3 Mechanical effects of MGO-mediated collagen crosslinking in demineralized and bonded dentin

Collagen crosslinking is known to enhance the mechanical stability of collagen-based tissues at both the nano- and macroscale. Glycation with methylglyoxal produces advanced glycation end products (AGEs), predominantly MG-H1, which accounts for nearly 90% of MGO-derived crosslinks. These intermolecular crosslinks stabilize the tropocollagen network, increasing resistance to mechanical deformation and limiting enzymatic access to cleavage sites. To evaluate the contribution of MGO-induced crosslinking to dentin durability, both the elastic modulus of demineralized dentin and the microtensile bond strength of resin-bonded dentin were assessed before and after collagenase challenge.

#### 3.3.1 Apparent elastic modulus of demineralized dentin

After 24 hours, untreated dentin exhibited an AEM of 1.07 ± 0.47 MPa. Glycation with 0.5 M and 1 M MGO produced modest, non-significant increases in stiffness (1.88 ± 0.44 MPa and 1.82 ± 0.48 MPa, respectively; p > 0.05). In contrast, 3 M MGO resulted in a pronounced and significant four-fold increase in AEM (4.01 ± 0.97 MPa; p < 0.0001). Following collagenase exposure, all groups showed reduced stiffness, reflecting enzymatic degradation. The decrease was most substantial in the untreated control, which dropped to 0.34 ± 0.38 MPa (p = 0.002). MGO-treated samples retained significantly higher AEM values (1.38–3.84 MPa; p < 0.05), with the 3 M group maintaining the greatest resistance to collagenase.

The dose-dependent increase in stiffness following MGO treatment aligns with previous findings showing that MGO-induced AGEs markedly enhance the nanoscale and bulk stiffness of collagen scaffolds [22, 36]. Riboflavin-mediated crosslinking produces similar glycation-like stiffening effects in demineralized dentin when photoactivated [37, 38], further supporting the role of crosslinks in reinforcing collagen architecture. The preserved AEM after collagenase challenge demonstrates that MGO effectively shields collagen fibrils from enzymatic degradation by limiting access to cleavage sites. These mechanical benefits are essential for improving the functional stability of dentinal substrates prior to adhesive bonding.

#### 3.3.2 Microtensile bond strength of resin-bonded dentin

Resin composite restorations achieve adhesion to dentin through the formation of a hybrid layer created by resin infiltration into the demineralized collagen matrix. To ensure that MGO treatment did not interfere with adhesive penetration, resin tag infiltration was evaluated using Rhodamine B-doped adhesive and confocal microscopy (Figure 5.A). No statistically significant differences were observed between the control and MGO-treated groups (ctrl: 30.3 ± 4.9 µm, 0.5 M MGO: 32 ± 3.1 µm, 1 M MGO: 26 ± 7.3 µm, 3 M MGO: 28.6 ± 6.8 µm; p > 0.05), indicating that MGO-induced glycation did not impair resin infiltration or hybrid layer formation.

**Figure 5.**
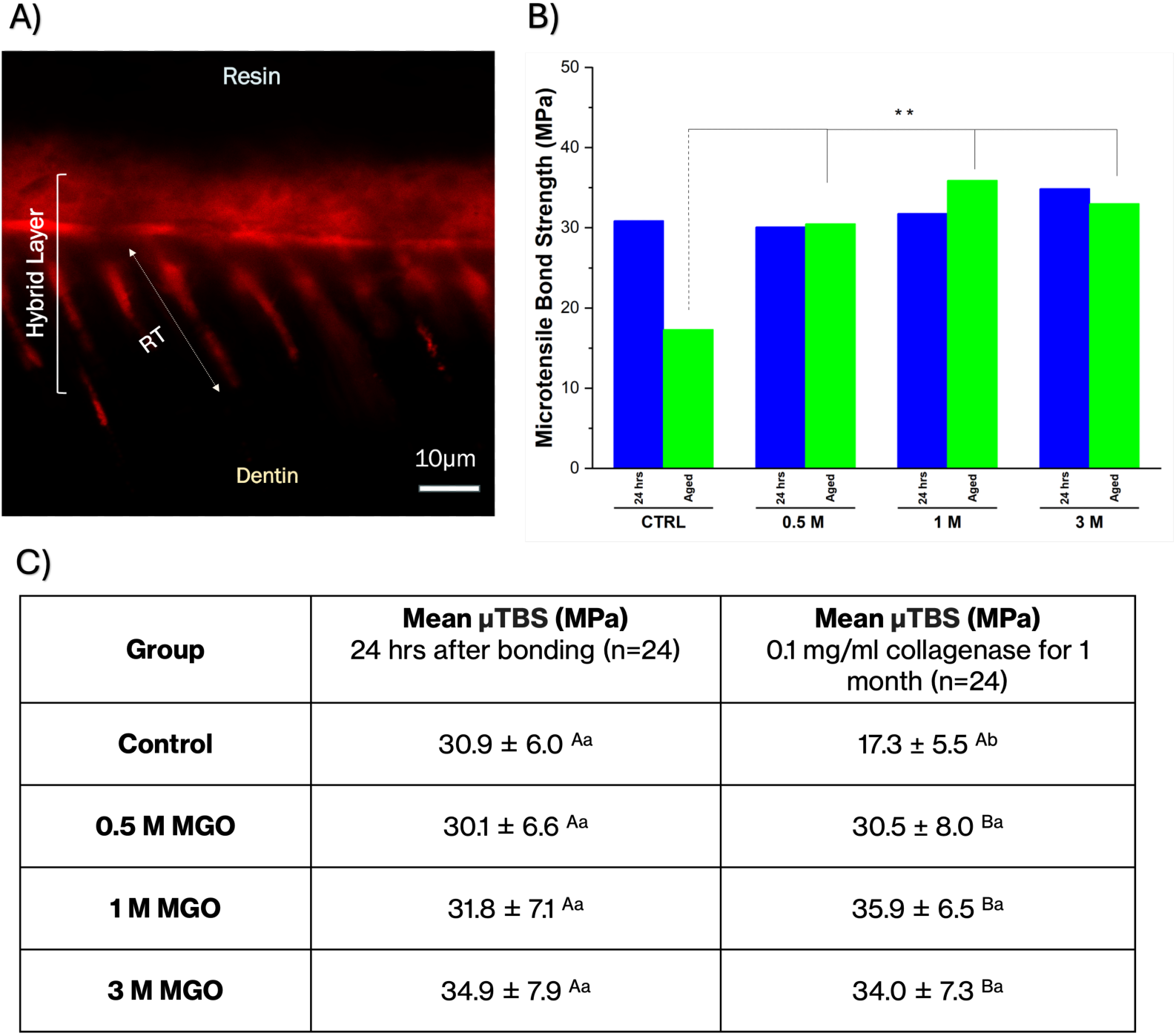
Microtensile bond strength (µTBS) of resin–dentin interfaces following MGO treatment. Bar graph and table showing mean µTBS values (± SD) for control and MGO-treated dentin (0.5 M, 1 M, and 3 M) measured 24 hours after bonding and after one month of incubation in 0.1 mg/mL collagenase (n = 24 beams per group). Lowercase superscript letters indicate statistically significant differences within each column (p < 0.05), and uppercase superscript letters indicate statistically significant differences within each row (p < 0.05).

Figure 5B and 5C summarize the effects of MGO treatment on resin–dentin bond strength at 24 hours and after enzymatic aging. At 24 hours, µTBS values for all MGO-treated groups (0.5–3 M) were statistically similar to the control (≈30–35 MPa), suggesting comparable early bonding performance. After one month of storage in 0.1 mg/mL collagenase, the control group showed a significant decline in bond strength (17.3 ± 5.5 MPa; p < 0.01). In contrast, MGO-treated specimens maintained markedly higher µTBS values (30.5–35.9 MPa), approximately double the degraded control (p < 0.01).

The preservation of bond strength in MGO-treated specimens supports the formation of protective collagen crosslinks within the hybrid layer, thereby restricting enzymatic access and preventing structural collapse during aging. While MGO does not influence immediate bond performance, it significantly enhances long-term interface durability. By stabilizing the collagen matrix, MGO reduces enzymatic degradation pathways that typically compromise resin-dentin bonds over time, underscoring its potential as a biomodification strategy for extending the longevity of adhesive restorations.

The preservation of bond strength in MGO-treated specimens supports the formation of protective collagen crosslinks within the hybrid layer, thereby restricting enzymatic access and limiting structural degradation during aging. While MGO did not influence immediate bond performance, it significantly enhanced long-term interface durability. These findings are consistent with a recent systematic review demonstrating that collagen crosslinking agents improve the long-term bond strength of resin–dentin interfaces by stabilizing the collagen matrix and reducing enzymatic degradation[14]. By reinforcing the collagen network, MGO appears to attenuate degradation pathways that typically compromise resin–dentin bonds over time, underscoring its potential as a biomodification strategy for extending the longevity of adhesive restorations.

##### The potential of MGO-glycation in restorative dentistry

Several studies have demonstrated the efficacy of external crosslinking agents in stabilizing collagen within the hybrid layer, mitigating enzymatic degradation, and strengthening resin-dentin bonds. The most common application of cross-linkers is as a pretreatment agent after acid etching, with considerable potential to enhance bond longevity[39]. Riboflavin is a widely tested glycation-like cross-linker known for forming covalent bonds with proline and lysine residues upon UV light activation. This process stabilizes the collagen matrix and inhibits matrix metalloproteinase (MMP) activity[39]. A study by Chiang et al. (2013) demonstrated that a 0.1% riboflavin solution effectively maintained high bond strength while minimizing nano-leakage[38]. Another agent, proanthocyanidin (PA), is a natural crosslinker with antibacterial, antioxidant, and anti-inflammatory properties that showed promising results in improving both immediate and aged bond strength[40, 41]. While MGO has been investigated mainly in tissue engineering of collagen scaffolds, limited research is available on its potential use in restorative dentistry applications.

MGO demonstrates significant potential as a collagen crosslinker for use as a primer applied to etched dentin before bonding to resin restorations, especially in cases where minimal invasive caries treatment (MICT) is required. MICT focuses on preserving pulp vitality while effectively managing carious lesions (with or without indirect pulp capping)[42, 43]. This approach is based on the understanding that carious dentin consists of two distinct layers: an outer layer of infected dentin, which is soft, highly porous, and densely populated with microorganisms; and an inner layer of affected dentin, which is firmer, less microbial, and retains the potential for remineralization[42]. In the partial caries removal technique, the infected dentin is excavated while the affected dentin is preserved to minimize the risk of pulp exposure. However, the affected dentin often displays hypomineralization, increased porosity, and disorganized collagen fibers, leading to compromised mechanical properties[43, 44]. Research has demonstrated that the caries-affected dentin-resin interface is highly prone to degradation, mainly due to elevated proteolytic activity (e.g., MMPs) and an increased likelihood of phase separation[44]. This is where MGO shows significant promise. In addition to its crosslinking ability, particularly when used in high concentrations, it exhibits potent antimicrobial and cytotoxic activity by inducing bacterial cell death through single-strand DNA breaks and DNA-protein cross-linking[45]. During cavity preparation for resin-based restorations, the application of MGO as a primer following acid etching is proposed as a potential strategy to sanitize the cavity prior to bonding, leveraging its antimicrobial properties[45]. It thoroughly sanitizes the remaining dentin by eliminating residual microorganisms that survived the etching process. Beyond its antimicrobial efficacy, MGO chemically interacts with the collagen matrix, forming intermolecular cross-links that functionalize dentinal collagen[22]. This enhances the matrix’s resistance to enzymatic degradation and improves its mechanical properties, potentially improving the hybrid layer while reducing the risk of secondary caries and bond failure. In endodontics, a 2022 study addressed the weakening of roots in endodontically-treated teeth, primarily caused by sodium hypochlorite irrigation[25]. The study investigated MGO as a protective agent, resulting in a significant increase in fracture resistance and a decrease in fracture depth[25]. Therefore, MGO emerges as a promising therapeutic agent for adhesive-based restorations by stabilizing the dentinal collagen matrix and enhancing its mechanical integrity and resistance to enzymatic degradation. In addition, MGO-treated dentin maintained resin–dentin bond strength under prolonged enzymatic challenge, indicating adequate protection of the hybrid layer against proteolytic degradation. It is worth noting that, like other cytotoxic materials used in dentistry (e.g., resin adhesive monomers or sodium hypochlorite)[46, 47], MGO requires careful handling. Proper isolation, such as using a rubber dam and controlled application with a micro-brush, can help minimize unintended effects during its use.

## 4 Conclusion

The use of methylglyoxal (MGO) before adhesive application represents a promising biomodification strategy to enhance the structural integrity of the resin–dentin interface. By selectively stabilizing the collagen matrix without affecting surface wettability, MGO crosslinking addresses a critical factor contributing to the limited longevity of resin-based restorations. These findings support the development of MGO as a collagen-targeted primer for improving long-term clinical outcomes.

## 5 Authors contributions

**Wafa Alzubaidi**, Conceptualization, Investigation, Methodology, Formal analysis, Writing – original draft, Writing –review & editing. **Bowen Wang, Steven Thorpe,** Investigation. **Mina Vaez, Mehrnoosh Neshatian**, Formal Analysis, Writing –review & editing. **Sebastian Aguayo,** Writing –review & editing. **Eszter Somogyi-Ganss**, Conceptualization, Writing –original draft, Writing –review & editing. **Laurent Bozec**, Conceptualization, Visualization, Funding acquisition, Methodology, Formal analysis, Writing –original draft, Writing –review & editing.

## 6. Acknowledgments

The authors express their gratitude to Natural Sciences and Engineering Research Council of Canada (NSERC), Ontario Research Fund (ORF), the Faculty of Dentistry at the University of Toronto and the Saudi Arabian Cultural Bureau for supporting the research. Acknowledgment is also extended to the CAMiLoD Imaging Facility at the Faculty of Dentistry for providing access to their equipment.

## 7 Conflict of Interest

The authors confirm that they have no financial or professional conflicts of interest that could have influenced the outcomes of this study.

## 8 Research funding

Natural Sciences and Engineering Research Council of Canada. Grant Numbers: RGPIN-2021-512186, and DH-2023-00130; Ontario Research Fund (ORF). Grant number: RE010-068; Joint Canadian Fund for Innovation & Ontario Research Fund – Small Infrastructure Fund #42809.

## Notes

### Competing Interest Statement

The authors have declared no competing interest.

